# Interleukin-36 upregulates type-I interferon responses in systemic lupus erythematosus by promoting the accumulation of self-nucleic acids

**DOI:** 10.1101/2025.10.20.683168

**Authors:** Emma J Welsh, Daniel McCluskey, Patrick Baum, Myles J Lewis, Francesca Capon

## Abstract

**Introduction:** Several studies have reported an up-regulation of interleukin (IL)-36 in the serum of patients with systemic lupus erythematosus (SLE). Here, we sought to define the mechanisms whereby IL-36 may contribute to the over-activation of type I Interferon (IFN) responses observed in SLE.

**Methods:** We carried out single-cell (sc)RNA-seq in healthy peripheral blood mononuclear cells treated with IL-36 (n=5 donors). We compared the genes and transcriptional networks that were induced by IL-36 with those that were upregulated in a published SLE scRNA-seq dataset (n=33 cases and 11 controls). In follow-up studies, we validated the effects of IL-36 on monocytes by real-time PCR (n=9 donors) and flow-cytometry (n=6).

**Results:** Classical monocytes were the immune population most affected by IL-36 treatment (n=203 Differentially Expressed Genes). In these cells, IL-36 upregulated transcriptional networks (regulons) driven by IRF7, a key activator of type I IFN responses. A similar upregulation of IRF7 regulons was observed in the monocytes of SLE cases, where measurements of IL-36 and IRF7 activity were significantly correlated (r=0.35, P=0.02). Experimental follow-up studies in human monocytes showed that IL-36 downregulates multiple RNAse genes (*RNASE1, RNASE6, RNASET2*). IL-36 treatment of monocytes also increased the percentage of apoptotic cells (45% vs 37% in untreated cells; P=0.001), which are a critical source of self-nucleic acids.

**Conclusion:** We find that IL-36 promotes monocyte apoptosis while downregulating self-nucleic acid clearance. Thus, IL-36 contributes to the accumulation of self-nucleic acids, a key driver of type I IFN responses in SLE.

## 1 Introduction

Systemic lupus erythematosus (SLE) is an autoimmune disorder associated with significant morbidity and mortality. The disease affects multiple organs, reflecting the damage caused by the deposition of autoantigen-autoantibody complexes(1). These contain a variety of nuclear components, including nucleic acids released from apoptotic cells. In fact, immune complexes containing self-DNA or self-RNA can trigger a potent type I interferon (IFN) response, which is a hallmark of SLE(1).

The pathological importance of type I IFN signaling was underscored by the study of monogenic forms of SLE and the genetic analysis of interferonopathies, a group of lupus-like diseases characterized by the upregulation of type I IFN responses(2). While genetically heterogeneous, these conditions are often caused by mutations that impair the effective clearance of self-nucleic acids. These disease alleles affect DNAse and RNAse genes, leading to the accumulation of self-DNA/RNA, with consequent activation of type I IFN responses(2). While the study of these rare conditions has provided valuable insights into pathogenic processes, less is known about the mechanisms deregulating nucleic acid clearance in multifactorial forms of SLE.

Interleukin-36α, -β and -γ (IL-36) are a group of closely related IL-1 family cytokines, which signal through the same receptor (IL-36R). Excessive IL-36 activity is a key driver of skin and systemic inflammation, as demonstrated in psoriasis and other immune-mediated skin diseases(3). Given this pro-inflammatory role, IL-36 has also been investigated in SLE. Several groups have reported that IL-36 levels are increased in the serum of affected individuals, where IL-36 concentrations correlate with disease activity(4, 5). We and others have also shown that IL-36 can upregulate type I IFN production in human plasmacytoid dendritic cells and type I IFN responsiveness in mouse keratinocytes(6, 7). These findings, however, were documented in cells exposed to viruses or viral mimics. Thus, the mechanisms whereby IL-36 may enhance type I IFN signaling in SLE have not been investigated. Here, we show that IL-36 promotes the accumulation of self-nucleic acids, which is a key driver of type IFN production in SLE.

## 2 Methods

### 2.1 Healthy Donor Recruitment

This study was undertaken according to the principles of the Declaration of Helsinki. Healthy donor blood samples were obtained from volunteers recruited at St John’s Institute of Dermatology (London Bridge research ethics committee approval: 16/LO/2190) or from leukocyte-enriched cones (NHS Blood and Transplant Service, Tooting, UK), depending on the application (see below). All participants granted their written informed consent.

### 2.2 Cell culture and IL-36 stimulation

Peripheral Blood Mononuclear Cells (PBMCs) were isolated by density gradient centrifugation of blood samples (for scRNA-seq and real-time PCR) or leukocyte cones (for flow cytometry). Cells were cultured in RPMI 1640 (Gibco) with 10% FBS (Gibco) and 1% Penicillin/Streptomycin (ThermoFisher). 10^6^ cells were treated with IL-36γ (50ng/ml) or PBS for 7h. Cells were then resuspended in PBS for flow-cytometry or stored in freezing medium for scRNA-seq.

Monocytes were purified from PBMCs using a human pan-monocyte isolation kit (Miltenyi Biotech). Cells were cultured in RPMI 1640 supplemented with 1% Penicillin/Streptomycin, 10% FBS, 1X MEM Non-Essential Amino Acids (Gibco) and 1M HEPES Buffer Solution (Gibco). 10^6^ cells were treated with IL-36γ (100ng/ml) or medium for 7h. Cell pellets were then frozen for RNA isolation.

### 2.3 scRNA-seq

PBMCs were processed using the Chromium Single Cell 3’ Reagent Kit v3 (10x Genomics). Pooled libraries were sequenced on an Illumina HiSeq4000 instrument and reads were processed with Cell Ranger v3.0.2 (10x Genomics). scRNA-seq data was analyzed using Seurat v4.0.4. Cells with mitochondrial gene content >20% and cells expressing <300 or >5000 genes were excluded. The presence of doublets was also confirmed using DoubletFinder. Datasets were normalized using LogNormalise with a 10^4^ scale factor.

Cell clustering was implemented with the FindNeighbours and FindClusters functions, using Clustree to identify the most appropriate resolution. Clusters were visualized with Uniform Manifold Approximation and Projection (UMAP) plots and cell identities were annotated based on the expression of key markers.

### 2.4 Analysis of differentially expressed genes and transcriptional networks

Differentially expressed genes (DEG) were identified using the Seurat FindMarkers function, based on a log(FoldChange)> log(1.2), FDR<0.05 and expression in >10% of examined cells. IL-36 scores were calculated in each affected individual as the median normalized expression of the top nine genes induced by IL-36 stimulation of healthy classical monocytes (Supplementary Table S1). Regulatory networks were inferred with SCENIC v1.2.4. Regulon activity was calculated for each cell and then binarized.

### 2.5 Generation of diffusion maps and pseudotime analysis

The Destiny package (v3.8.0) was used to visualize binary regulon activity matrices in a three-dimensional space. The slingshot package (v2.2.1) was used to analyze the binary regulon activity matrices generated by SCENIC. Cells were annotated as untreated or treated, cell clusters were created using WhichCells and added to the Seurat object metadata to highlight regulons of interest.

### 2.6 Real-time PCR and apoptosis assay

Following RNA isolation and cDNA synthesis, gene expression was measured by real-time PCR, using the primers reported in Supplementary Table S2. Transcript levels were normalized against *GAPDH*.

The abundance of apoptotic monocytes was determined by staining PBMCs with propidium iodide (PI, from ApoDETECT Annexin V-FITC kit, Invitrogen) and with antibodies against CD14, CD16 and Annexin V (Supplementary Table S3). Cells were acquired on a CytoFLEX LX analyser (Beckman Coulter Life Sciences) and analyzed with FlowJo v10.8.1 (Beckton, Dickson & Company).

### 2.7 Statistical Tests

Two-tailed statistical tests were implemented in RStudio v1.4.1 or GraphPad Prism v9. Differences between two groups were assessed with the Mann-Whitney or Wilcoxon signed-rank test, as appropriate. The overlap between DEG detected in different scRNA-seq datasets was assessed with RStudio GeneOverlap function. Odds ratios over genomic background were computed using Fisher’s exact test. For the analysis of SCENIC output, statistical significance was assessed with a chi-squared test and p-values were adjusted using the Bonferroni correction.

## 3 Results

### 3.1 Single-cell analysis of IL-36 treated PBMCs

To investigate the leukocyte immune pathways that are activated by IL-36, we carried out scRNA-seq in PBMCs that had been exposed to the cytokine. To avoid the confounding effects of end-stage inflammation, we carried out the experiment in the cells of healthy donors (n=5). After quality control, we obtained data from 51,691 PBMCs (Supplementary Figure 1A-B), which we separated into ten clusters recapitulating the main leukocyte subsets (Figure 1A-B). As expected for a short stimulation, there was no difference in cell population abundance between treated and untreated samples (Supplementary Figure 1C). Conversely, IL-36 exposure showed a variable impact on transcript levels, with the number of differentially expressed genes (DEG) varying by two orders of magnitude across cell populations (Figure 1C). As the highest numbers of DEG were observed in classical monocytes (n=480), intermediate/non-classical monocytes (n=193) and NK cells (n=31) (Figure 1D-E, Supplementary Figure 1D), we focused our follow-up studies on these populations.

**Figure 1:**
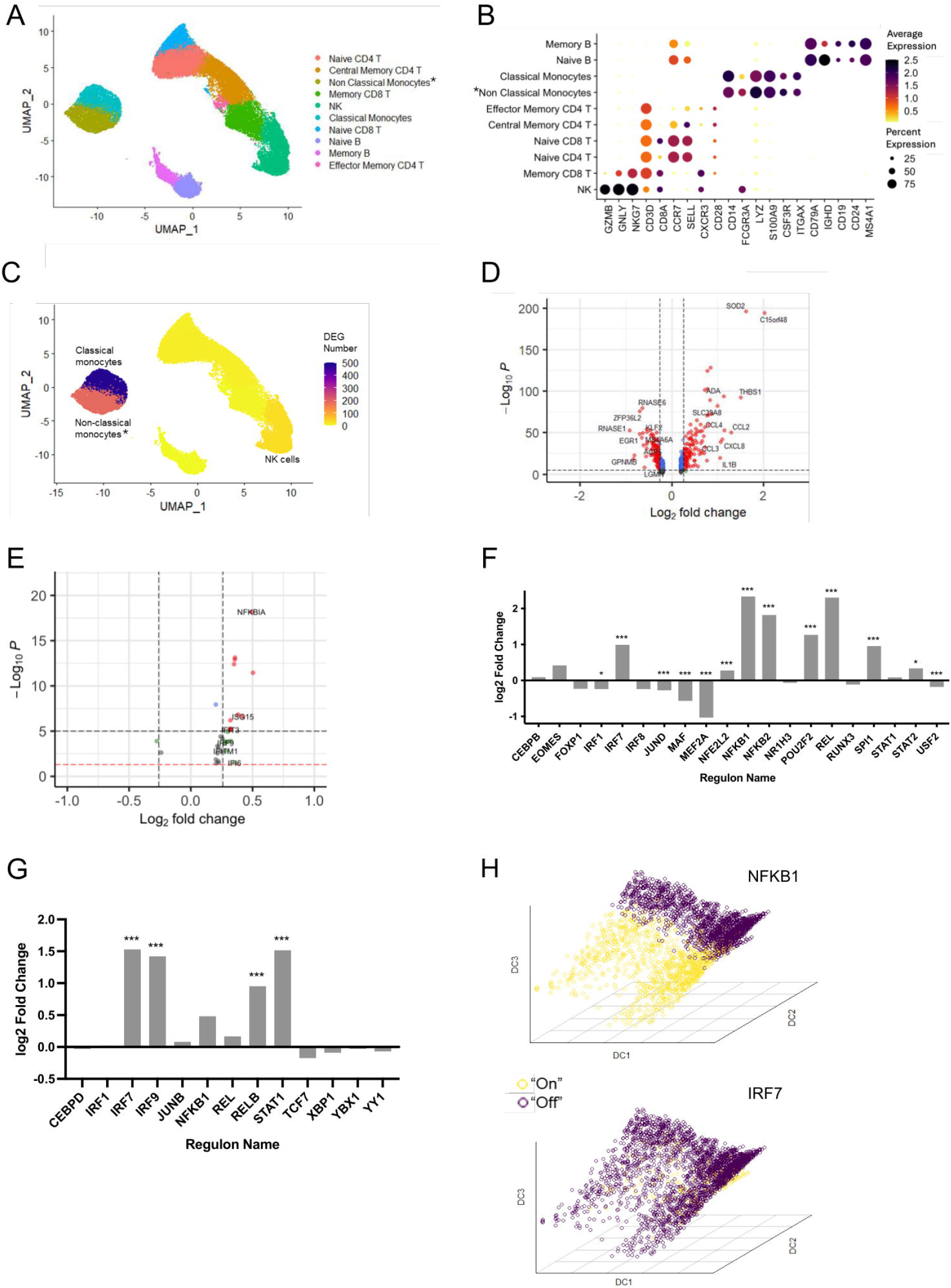
Single-cell RNA-seq analysis of healthy PBMC treated with IL-36. (A) UMAP of 51,691 single cells visualized as annotated clusters; *population including non-classical and intermediate monocytes. (B) Dot plot showing the expression of key markers used for cluster annotation. (C) UMAP visualization of the number of differentially expressed gene (DEG) detected in each cluster, following IL-36 treatment; (D-E) Volcano plots showing the DEG detected in classical monocytes (D) and natural killer (NK) cells (E). Horizontal and vertical dashed lines show the thresholds for statistical significance and fold change, respectively. Genes yielding a significant p-value or log2foldchange>log1.2 are represented by blue and green dots respectively; red dots indicate both thresholds were met. (F-G) Bar plots summarizing the changes in activity for the 20 regulons detected in classical monocytes (F) and the 13 regulons detected in NK cells (G). Positive log2foldchanges indicate increased activity in samples treated with IL-36. \**P*<0.05; \*\*\**P*<0.001 (χ^2^ test followed by Bonferroni correction). (H) Three-dimensional diffusion maps of NFKB1 and IRF7 activation in classical monocytes. Colors show cells where the regulon is active (yellow) or inactive (purple). DC, diffusion component.

### 3.2 IL-36 treatment activates interferon-regulated transcriptional networks

To investigate the effects of IL-36 treatment, we used SCENIC to infer the activation of transcriptional networks in the selected cell populations. In classical monocytes, this identified 20 regulons (sets of co-expressed genes sharing a binding site for the same transcription factor). When their activity (quantified as number of cells expressing the regulon) was compared in stimulated vs unstimulated samples, statistically significant differences were observed for regulons driven by NFKB1, REL and NFKB2 (log2FC≥1.8, FDR<0.001 for all) (Figure 1F). This is in line with the well-documented effects of IL-36 on NF-κB signaling(8). Surprisingly, however, we also observed that IL-36 treatment promoted the activation of IRF7 (log2FC=0.99, FDR<0.001), a key upstream regulator of type I IFN production (Figure 1F). A similar, although less conspicuous, activation pattern was observed in intermediate/non-classical monocytes, where IL-36 treatment induced regulons related to NF-κB signaling (NFKB1, log2FC:1.1, FDR<0.001) and type I IFN responses (IRF7, log2FC:0.23, FDR<0.01; STAT1, log2FC:0.57, FDR<0.001) (Supplementary Figure 1E). Finally, the analysis of NK cells revealed a robust activation of IRF7 (log2FC:1.5), STAT1 (log2FC:1.5) and its binding partner IRF9 (log2FC:1.4) (FDR<0.001 for all) (Figure 1G), in cells from IL-36 treated samples. While *IFNA* and *IFNB1* transcripts were undetectable across the entire dataset, the upregulation of interferon signature genes (*ISG15, IFIT3*) in NK cells, further confirmed the strong activation of type I IFN responses in this population (Figure 1E).

Taken together, these observations suggest that IL-36 promotes the activation of IRF7 in classical monocytes, leading to the transcription of type I IFN genes. These are likely to signal through STAT1 and IRF9 to upregulate interferon stimulated genes (ISG) and IRF7 regulons in NK cells (and to a lesser extent in intermediate/non-classical monocytes).

To further investigate this possibility, we visualized the classical monocyte data using diffusion mapping, a dimensionality reduction technique that preserves the pseudo-temporal ordering of cells(9). Using the Destiny package, we showed that the NFKB1 and IRF7 regulons are active in distinct cell neighborhoods (Figure 1H). Accordingly, trajectory analysis confirmed that the two regulons are upregulated at different points in pseudotime (Supplementary Figure 1F). These observations indicate that IL-36 induces a sequential rather than parallel activation of the NFKB1 and IRF7 responses.

### 3.3 SLE monocytes display a signature of IL-36 activation

To investigate the disease relevance of our findings, we retrieved an scRNA-seq dataset generated in the PBMCs of 33 SLE cases and 11 healthy donors(10). Following quality control, cell clustering and annotation (Figure 2A-B), we extracted 42,080 classical monocytes and compared gene expression in cases vs controls. This identified 320 DEG, of which 203 were upregulated (Figure 2C). When the latter were compared to the 216 monocyte genes that were induced by IL-36 treatment of healthy PBMCs, a substantial overlap was observed (n=29 shared genes; *P*=1.24×10^-5^; odds ratio over genomic background: 2.76) (Figure 2D). Thus, IL-36 signature genes are over-represented among the DEG detected in SLE classical monocytes.

**Figure 2:**
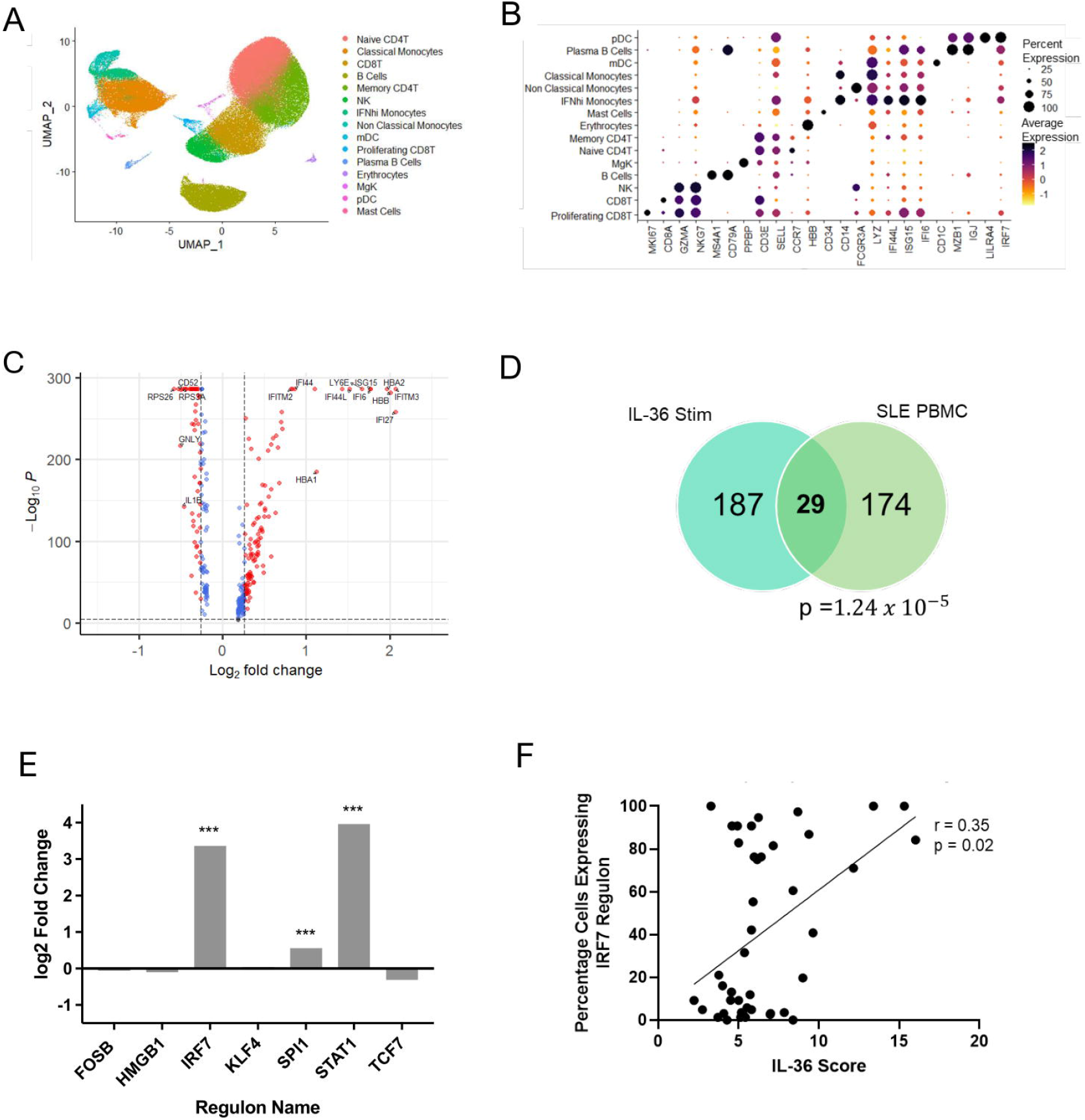
Identification of an IL-36 signature in SLE PBMCs. (A) UMAP of 280,725 single cells visualized as annotated clusters. (B) Dot plot showing the expression of key markers used for cluster annotation (C) Volcano plot showing the DEG detected by comparing the classical monocytes of SLE cases with those of healthy controls. Horizontal and vertical dashed lines show the thresholds for statistical significance (FDR<0.05) and fold change (log2foldchange>log1.2), respectively. Differentially expressed genes meeting both thresholds are represented by red dots, those that only meet the significance threshold are shown as blue dots. (D) Overlap between the genes induced by IL-36 stimulation and those up-regulated in SLE classical monocytes. A hyper-geometric test was used to determine statistical significance. (E) Bar plot summarizing the changes in activity for the seven regulons detected in classical monocytes. Positive log2foldchange indicates increased activity in SLE samples compared with healthy donors. \*\*\**P*<0.001 (χ^2^ test followed by Bonferroni correction). (F) Correlation between IL-36 scores and percentage of cells expressing the IRF7 regulon in the same individual (n=33 SLE cases). Statistical significance was assessed with Spearman rank correlation test.

We next examined the activation of transcriptional networks. As expected, we observed that STAT1 and IRF7 regulons were significantly upregulated in the classical monocytes of SLE cases vs those of healthy controls (log2FC: 3.96 and 3.36, respectively; FDR<0.001) (Figure 2E). We then focused on the patient group, measuring IL-36 and IRF7 activation in each affected individual. We found a significant correlation between IL-36 scores (calculated as the median expression of multiple IL-36 signature genes, see Methods) and percentage of cells with active IRF7 regulons (r=0.35, *P*=0.02) (Figure 2F). Thus, the upregulation of IL-36 signaling in classical monocytes of SLE patients correlates with the abnormal activation of type I IFN responses.

### 3.4 IL-36 affects RNAse gene expression and apoptosis in SLE monocytes

To investigate the mechanisms whereby IL-36 may upregulate type I IFN responses, we examined the DEG detected in IL-36 stimulated PBMCs. Interestingly, we found that the expression of *IFNAR1* and *IFNAR2* (encoding the IFNα/β receptor) was unaffected by cytokine treatment. Conversely, we observed that *RNASE1* was the most downregulated gene in classical monocytes (log2FC: -0.9; FDR<10^-50^) and that the expression of other enzymes promoting nucleic acid degradation (*RNASE6, RNASET2, DNASE2, PLD3, SAMHD1*) was also reduced in these cells (Figure 3A). Of note, three of the downregulated genes have been associated with interferonopathies(2, 11) (Figure 3A).

**Figure 3:**
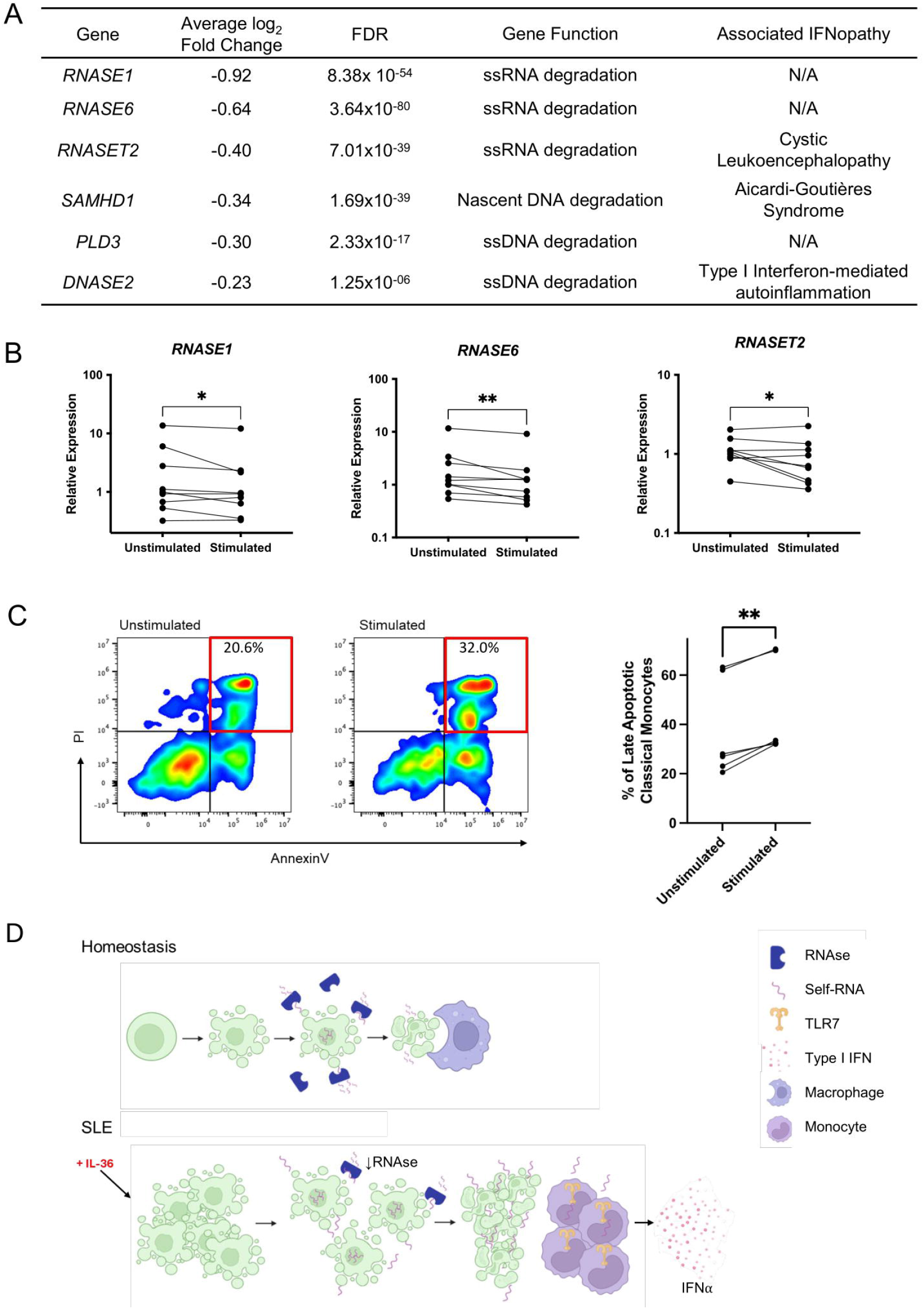
IL-36 downregulates RNAse gene expression and upregulates apoptosis in healthy monocytes. (A) RNAse and lupus-related genes downregulated in classical monocytes from IL-36 treated PBMCs. ss, single-stranded. (B) Paired dot plots showing relative gene expression in IL-36 stimulated monocytes (n=9 healthy donors). Expression levels were measured by real-time PCR and normalized to *GAPDH* mRNA levels. Each line represents one donor. \**P*<0.05; \*\**P*<0.01 (ratio paired t test). (C) Left: representative flow cytometry plots showing the abundance of late apoptotic classical monocytes in unstimulated vs. IL-36 stimulated cells. Classical monocytes were first gated as CD14+/CD16-cells. Late apoptotic cells were then identified as a PI+/Annexin+ population (red box). PI, propidium iodide. Right: paired dot plot showing the proportion of classical monocytes identified as late apoptotic, in unstimulated vs. IL-36 stimulated cells (n=6 donors). Each line represents one donor. \*\**P*<0.01 (paired t-test). (D) Proposed effect of IL-36 on classical monocytes. Under homeostatic conditions, nucleic acids released during apoptosis are degraded by RNAses and DNAses, while apoptotic cells are phagocytosed by macrophages. In SLE, circulating IL-36 upregulates apoptosis while downregulating RNAse expression. The resulting nucleic acid build-up leads to the activation of TLR7, amplifying Type I IFN responses. Image created with biorender.com

To validate our observations, we treated healthy monocytes (n=9 donors) with IL-36 and measured gene expression by real-time PCR, focusing on the three genes showing the strongest downregulation in the scRNA-seq dataset (*RNASE1, RNASE6, RNASET2*). We found that in all cases, IL-36 caused a modest but reproducible reduction in gene expression (fold-change: 0.75-0.83; *P*<0.05) (Figure 3B).

Further analysis of the DEG detected in IL-36 treated PBMCs revealed that two of the most upregulated genes in classical monocytes (*C15orf48, THBS1;* log2FC>1.5; FDR<10^-90^ for both) contributed to apoptosis(12, 13). The expression of other proapoptotic genes was also elevated in these cells (*BID, CASP1, CASP4*; FDR<10^-6^ for all) (Table S4). To follow up these findings, we examined the effects of IL-36 on monocyte apoptosis. We found that cytokine treatment significantly increased the abundance of late apoptotic cells among classical monocytes (45.2% vs 37.3% at baseline; *P*=0.001; n=6 donors) (Figure 3C).

These observations suggest that IL-36 upregulates nucleic acid accumulation in classical monocytes, by promoting apoptosis while reducing self-DNA and self-RNA clearance.

## 4. Discussion

The aim of our study was to elucidate the mechanisms whereby IL-36 contributes to the activation of type I IFN responses in SLE. We first investigated the effects of the cytokine through the unbiased scRNA-seq analysis of IL-36 treated PBMCs. By examining the number of DEG induced by cytokine exposure, we identified classical monocytes as the main IL-36 responders in the circulation. This is in keeping with the robust expression of the IL-36 receptor on the surface of these cells(6).

To further explore our gene expression findings, we also analyzed transcriptional networks. We found that classical monocytes exposed to IL-36 increase the activity of NFKB1 and IRF7 regulons. The latter are also upregulated in SLE monocytes, where we observed a significant correlation between IL-36 and IRF7 activation. Of note, IRF7 acts downstream of innate immune pathways (e.g. TLR3/TLR7/TLR9 signaling) that can be triggered by endogenous nucleic acids. In this context, our findings suggest that IL-36 may influence the accumulation of self-DNA/RNA. This hypothesis is supported by the results of our follow-up experiments. We found that IL-36 promotes the apoptosis of classical monocytes, an observation that mirrors the pro-apoptotic effects of the cytokine in lung epithelial cells(14). We also showed that IL-36 downregulates three enzymes that degrade TLR7 ligands *(RNASE1, RNASE6, RNASET2)*. Thus, our findings suggest that IL-36 has a dual effect. It promotes apoptosis, causing the release of self-nucleic acids in the extra-cellular space and downregulates self-RNA clearance, leading to the activation of IRF7 (Figure 3D). Interestingly, IL-36 cytokines are upregulated by TLR signaling(8), which could further amplify inflammatory responses.

Our study has some limitations. The PBMC stimulation experiment was not sufficiently powered to detect DEG in rare leukocyte populations such as plasmacytoid dendritic cells (pDCs). Of note, pDCs are the main IFNα producers in the circulation and we previously showed that IL-36 can promote TLR9 translocation in these cells(6). Thus, further studies will be required to dissect the effects of IL-36 signaling in SLE pDCs.

We also acknowledge that the effects of IL-36 on RNAse expression were modest and not comparable to those of the loss-of-function mutations causing interferonopathies. At the same time, IL-36 reduced the levels of multiple nuclease genes, and we expect that the collective effect of these changes would be a self-nucleic acid build-up. Thus, our study identifies IL-36 as a cytokine that is likely to potentiate type I IFN inflammatory responses in SLE. Given the proven efficacy of IL36R blockers in severe psoriasis (generalized pustular psoriasis)(15), these findings warrant further mechanistic studies of IL-36 signaling in SLE.

## 5 Conflict of Interest

FC has received grant funding and consultancy fees from Boehringer-Ingelheim. PB is a Boehringer-Ingelheim employee.

## 6 Author Contributions

EJW: Formal Analysis, Writing – review & editing, Visualization, Investigation; DM: Formal Analysis, Writing – review & editing, Investigation; PB: Investigation; MJL: Supervision, Funding acquisition, Writing – review & editing; FC: Conceptualization; Supervision, Funding acquisition, Writing – original draft, Writing – review & editing

## 7 Funding

This research was supported by National Institute for Health Research (NIHR) Biomedical Research Centre based at Guy’s and St Thomas’ NHS Foundation Trust and King’s College London, and at Barts Health NHS Trust. EJW was funded by a Versus Arthritis PhD scholarship (grant 22506). DM was supported by the UK Medical Research Council (MRC, grant MR/R015643/1) and King’s College London as a member of the MRC Doctoral Training Partnership in Biomedical Sciences. FC is supported by the BC Leading Edge Endowment Fund. The views expressed in this publication are those of the authors and not necessarily those of the NHS or NIHR.

## 9 Data Availability Statement

The single-cell RNA-seq datasets generated for this study will be available in the Gene Expression Omnibus (GEO).

