## Supplementary Material for "Interleukin-36 upregulates type-I interferon responses in systemic lupus erythematosus by promoting the accumulation of self-nucleic acids"

### 1. Supplementary Figures

A

| Sample Group | Mean n. cells per sample (SD) | Mean n. reads per cell (SD) | Mean n. genes per cell (SD) |
|---|---|---|---|
| Unstimulated | 5,059 (681) | 6,231 (3644) | 1,904 (765) |
| Stimulated | 4,504 (488) | 6,311 (3796) | 1,904 (749) |

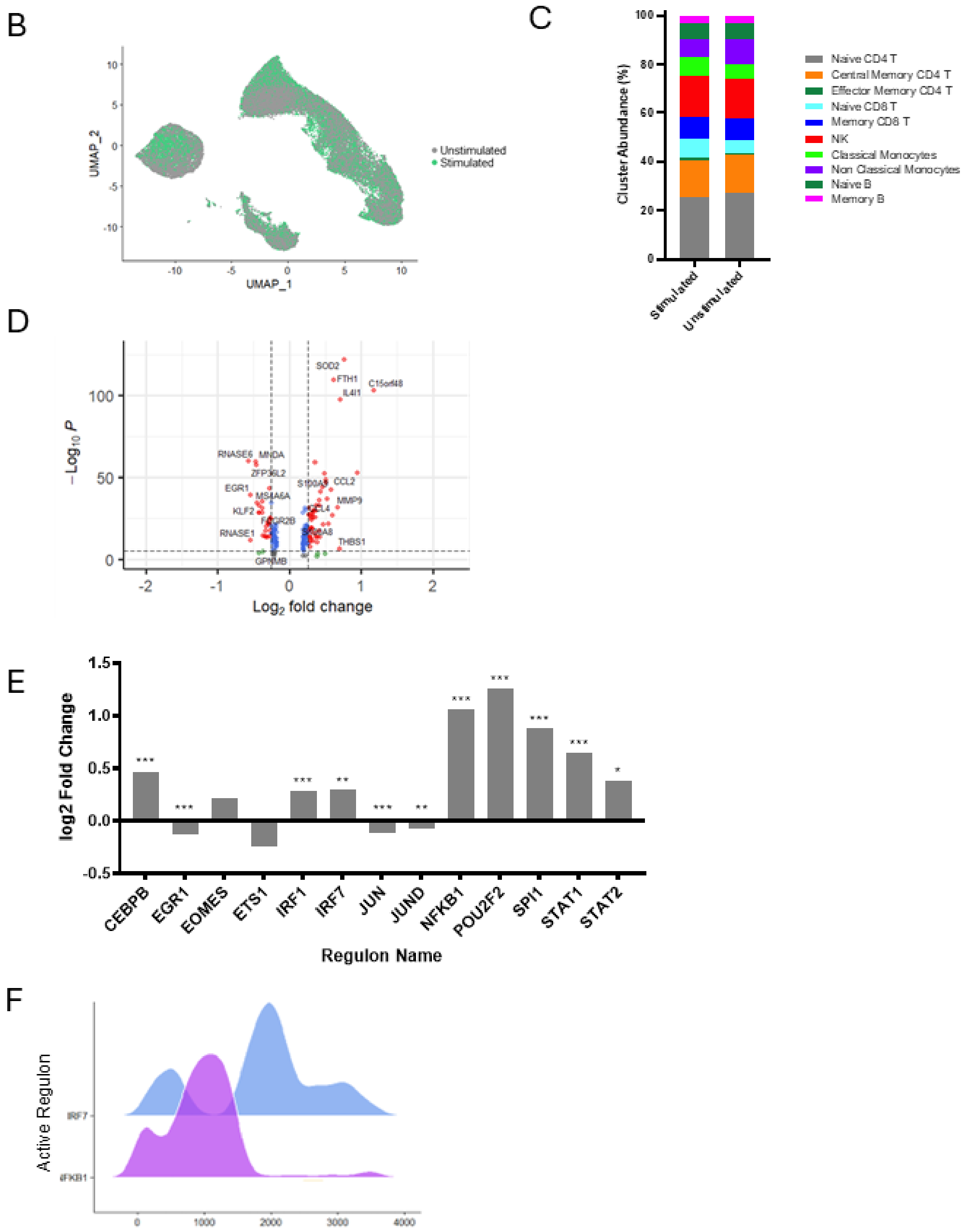


**Supplementary Figure S1**

(A) Single-cell RNA sequencing output for the IL-36 stimulation experiment. SD, standard deviation. (B) UMAP of 51,691 single cells visualized by treatment status. (C) Volcano plot showing the DEG detected in non-classical monocytes following IL-36 stimulation. Horizontal and vertical dashed lines show the thresholds for statistical significance (FDR<0.05) and fold change (log2foldchange>log1.2), respectively. Differentially expressed genes meeting both thresholds are represented by red dots, those that only meet the significance threshold are shown as blue dots. (D) Bar plots summarizing the changes in activity for the 13 regulons detected in non-classical monocytes. Positive log2foldchanges indicate increased activity in samples treated with IL-36. *$P$<0.05; **P<0.01; ***$P$<0.001 ($\chi^2$ test followed by Bonferroni correction). (E) Slingshot pseudotime analysis of classical monocytes displaying NFKB1 or IRF7 regulon activation.

A

| Sample Group | Mean n. cells per sample (SD) | Mean n. reads per cell (SD) | Mean n. genes per cell (SD) |
|---|---|---|---|
| Healthy Donor | 7774 (2074) | 3855 (1950) | 1099 (374) |
| SLE | 5916 (2377) | 3824 (2289) | 1137 (464) |

B

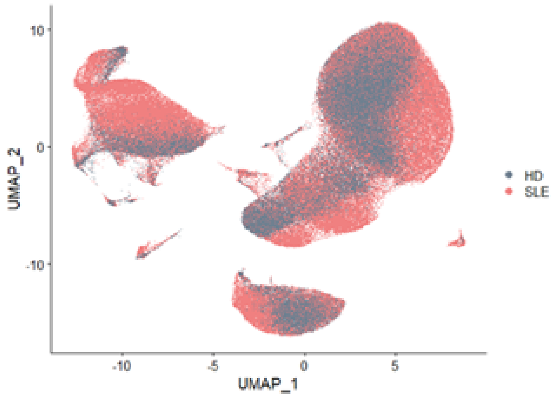


**Supplementary Figure S2:**

(A) Single-cell RNA sequencing output for the SLE case control dataset. SD, standard deviation. (B) UMAP of 280,725 single cells visualized by participant group.

### 2. Supplementary Tables

**Table S1:** Genes underlying the IL-36 score

| Gene Name | avg_log2FC | FDR |
|---|---|---|
| *C15orf48* | 2.01 | $6.2 \times 10^{-195}$ |
| *SOD2* | 1.62 | $9.2 \times 10^{-197}$ |
| *THBS1* | 1.50 | $7.6 \times 10^{-93}$ |
| *CCL2* | 1.29 | $3.5 \times 10^{-51}$ |
| *CCL4* | 1.14 | $1.3 \times 10^{-53}$ |
| *ADA* | 1.13 | $3.0 \times 10^{-94}$ |
| *CCL3* | 1.07 | $3.6 \times 10^{-39}$ |
| *IL1B* | 1.05 | $3.9 \times 10^{-20}$ |
| *SLC39A8* | 0.99 | $6.5 \times 10^{-83}$ |

FC; fold change, FDR; false discovery rate.

**Table S2:** Real-time PCR primers

| Target gene | Primer sequence (5' to 3') |
|---|---|
| *GAPDH* | F: CGGAGTCAACGGATTTGGTC |
| | R: AATGAAGGGGTCATTGATGGCA |
| *RNASE1* | F: AGCTGCAGATCCAGGCTTT |
| | R: CTGCCGCTGGAATTTCTTGG |
| *RNASE6* | F: GAGACCAGAAAAGATGGTGCT |
| | R: GAGCCTTGGTGAGACGCTTA |
| *RNASET2* | F: ATACATGGACTATGGCCCGA |
| | R: TGCGATTGGGAAACGAGTGA |

F, forward; R, reverse.

**Table S3:** Flow Cytometry Antibodies

| Fluorochrome | Target | Clone | Cat. No | Supplier | Dilution |
|---|---|---|---|---|---|
| AF700 | CD14 | 63D3 | 367114 | BioLegend | 1:100 |
| APC | CD16 | 3G8 | 302011 | BioLegend | 1:50 |
| FITC | Annexin V | N/A | 33-1200 | Thermo Fisher | N/A |

AF700, Alexa Fluor 700; APC, Allophycocyanin; FITC, Fluorescein isothiocyanate; N/A, not applicable.
